# The non-deterministic genotype-phenotype map of RNA secondary structure

**DOI:** 10.1101/2023.02.27.530309

**Authors:** Paula García-Galindo, Sebastian E. Ahnert, Nora S. Martin

## Abstract

Selection and variation are both key aspects in the evolutionary process. Previous research on the mapping between molecular sequence (genotype) and molecular fold (phenotype) has shown the presence of several structural properties in different biological contexts, implying that these might be universal in evolutionary spaces. The deterministic genotype-phenotype (GP) map that links short RNA sequences to minimum free energy secondary structures has been studied extensively because of its computational tractability and biologically realistic nature. However, this mapping ignores the phenotypic plasticity of RNA. We define a GP map that incorporates non-deterministic phenotypes, and take RNA as a case study; we use the Boltzmann probability distribution of folded structures and examine the structural properties of non-deterministic (ND) GP maps for RNA sequences of length 12 and coarse-grained RNA structures of length 30 (RNAshapes30). A framework is presented to study robustness, evolvability and neutral spaces in the non-deterministic map. This framework is validated by demonstrating close correspondence between the non-deterministic quantities and sample averages of their deterministic counterparts. When using the non-deterministic framework we observe the same structural properties as in the deterministic GP map, such as bias, negative correlation between genotypic robustness and evolvability, and positive correlation between phenotypic robustness and evolvability.

## I. INTRODUCTION

Biological information is stored in our genome in DNA and RNA as linear sequences composed of a nucleotide quaternary alphabet [1]. In nature, RNA is commonly found as a single stranded sequence that can fold, through nucleotide interactions, into a three-dimensional molecular structure. For short RNAs this three-dimensional structure can be coarse-grained to a two-dimensional representation of the fold (secondary structure) [2]. The biological functions that RNA executes in the cell depend on its folded structure [2], and so a direct link between RNA sequence, its minimum free energy (MFE) fold, and its function can be constructed [3, 4]. The sequence-structure mapping can be called the genotype-phenotype (GP) map for RNA, since the sequence is the biological information carrier (genotype) and the fold has a direct implication on function and therefore selection (phenotype) [4]. For RNA of length *L,* the GP map involves all possible L-sequences mapped to their respective MFE secondary structures, in which each sequence is connected to its point-mutation neighbours [5]. The GP map can then be interpreted as a road-map outlining the accessibility of phenotypes in terms of point mutations of genotypes [5]. Constructing this mapping is tractable for short RNAs with the currently available computational tools [6, 7], which is one of the main reasons why the GP map of RNA secondary structure has been studied so extensively [5].

The study of different GP maps which are not necessarily sequence-structure but represent a different phenotype (e.g. for proteins [8–10], metabolic networks [5] and GRNs [11]) has shown the prevalence of structural properties of the GP map that are common with the RNA sequence-structure map, suggesting that the GP maps exhibit properties that are characteristic of evolutionary spaces in general [1, 5]. These properties include redundancy, bias, correlations between evolvability (i.e. the potential for phenotypic changes through mutations) and robustness (i.e. the probability of phenotype-preserving mutations), and the presence of neutral spaces and genetic correlations [5]. First, most GP maps display redundancy, which means that there are many genotypes mapping to a single phenotype, for example many sequences folding into a single structure in the RNA GP map. The set of genotypes mapping to the same phenotype is called a neutral space [1], or a neutral network if they are connected by point mutations [1, 12]. Secondly, most GP maps are biased, which means that some phenotypes are much more redundant than others and thus have much larger neutral spaces. Thirdly, GP maps show a correlation between evolvability and robustness that is negative at genotypic level and positive at the phenotypic level [13]. Finally, GP maps usually display genetic correlations, which means that robustness is much higher than we would expect in a null model that simply accounts for neutral space sizes [14]. It is neutral spaces and networks in GP maps that explain the emergence of the more general concept of biological robustness [1, 13]. In our discussions so far, and in most of the GP map literature more widely, GP maps are deterministic many-to-one mappings, in which each genotype maps to one phenotype. For RNA, the minimum free energy (MFE) fold represents this deterministic phenotype [1, 5, 15]. However, in nature, phenotypes can be stochastic, either when the environment is variable, or even in a constant environment. For example, RNA folding is stochastic in a constant environment because of the inherent thermal fluctuations of the molecules at finite temperature [15]. Therefore, the nucleotide interactions create a range of possible folds described by the Boltzmann distribution. The structural phenotype of an RNA molecule is in fact plastic because it is fluctuating in time through Boltzmann-distributed suboptimal structures [15]. Seen in another way, at any one instant, a large set of RNAs of the same genotype, placed in an environment of temperature *T*, will exhibit a range of folded structures, where the amount of genotypes with fold *p* will be proportional to the Boltzmann probability *P*(*p*|*g*) = exp((*G_ens_* – *G_p_*)/*kT*), where *G_ens_* is the ensemble free energy (*G_ens_* = −*kT* · ln*Z* with partition function *Z*) and *G_p_* is the free energy that corresponds to structure *p* [15] (c.f. Fig.1). Meyers and Fontana’s highly influential work included plasticity in the RNA GP map [15]. They introduced an intrinsic feature of the plastic RNA GP map called plastogenetic congruence which is a correlation between a genotype’s suboptimal structures and the MFE structures of its point-mutation neighbours [15]. This GP map feature was also shown to create a negative correlation between plasticity and robustness [15]. The work from Meyers and Fontana as well as further more recent work [16–21] that includes plasticity, has improved our understanding of the plastic GP map but has not fully addressed how concepts such as robustness, evolvability and neutral spaces can be quantified for the non-deterministic case in a way that is consistent with averages over their deterministic counterparts.

**FIG. 1.**
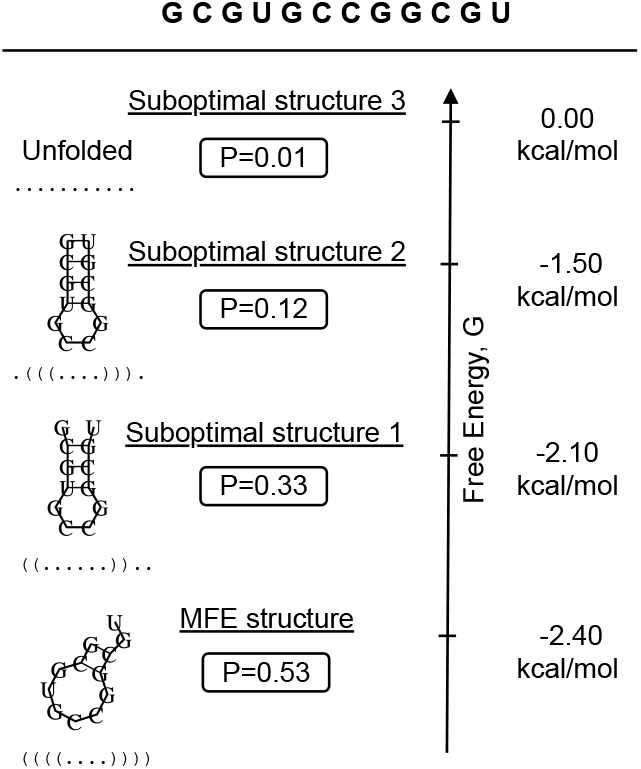
RNA plasticity example. Schematic of an example set of Boltzmann distributed suboptimal structures for genotype GCGUGCCGGCGU with their corresponding free energies G and normalised probabilities P. The unfolded case is included as a suboptimal structure with characteristic free energy G=0.0 kcal/mol. Secondary structures and energies are calculated using the ViennaRNA subopt command with default values [7].

Thermal fluctuations are ubiquitous in molecular environments, which means that many-to-many GP maps that include plasticity are more biologically realistic in the context of molecular evolution. These many-to-many maps can have nontrivial structural properties such as plastogenetic congruence and previous studies of plasticity suggest its possible impact on evolutionary dynamics and ecology, such as its ability to expedite evolution [15, 22–24] or facilitate the ecological coexistence of species [25, 26]. However, the extent to which plastic phenotypes impact evolution is still unclear [27, 28]. The strength of the GP map framework lies in its ability to clearly quantify the sequence-to-structure relationship [5]. The study of GP maps has advanced greatly due to the rapid increase in computational power witnessed over recent decades, as well as continual improvement of the tools used to study these high-dimensional spaces [28]. But also particularly important, has been the formulation of a general quantifying framework. Quantities such as robustness and evolvability have been key to the exploration of how different GP maps compare and the finding of GP map’s quasi-universal properties, which has eventually resulted in significant progress in understanding variation [5, 28]. So, accordingly, we expect that defining an analogous framework for the plastic GP map can be a baseline for further exploration of plastic GP maps in general, of any biological context, which could help bring answers on plasticity’s relationship to evolution.

The terms plasticity [15], promiscuity [28], intra-genotypic variability [29], non-determinism [5] and most recently, ‘probabilistic GP map (PrGP map)’ [20] have been used to describe many-to-many GP maps. Here we will use ‘non-deterministic (ND)’. The ND GP map is defined as a many-to-many mapping where the phenotypes of each genotype form a probability distribution, and for clarity we will refer to a general deterministic GP map, as ‘D GP map’. In the case of RNA, the MFE structure is usually taken to be the deterministic phenotype since it has the highest probability of folding and we will therefore compare our results from the full ND map to this MFE map. The work is structured as follows: first, we mathematically define non-deterministic quantities that are designed to correspond to averages of deterministic quantities over an ensemble. Then we validate this correspondence for both RNAs of *L* = 12 (conventional sequence-structure mapping) and *L* = 30 (RNAshapes [30, 31]). Following that, the deterministic quantities and their relationships are compared to the non-deterministic case to gain a deeper insight into the differences between the RNA ND GP map and the deterministic GP map. The computational methods used to construct the ND GP maps are explained at the end of this paper.

## II. RESULTS AND DISCUSSION

In order to generalise GP map measurements such as robustness, evolvability, and neutral space size to non-deterministic GP maps we take a stochastic approach. The stochastic versions of these measurements will quantify the *averages* of the corresponding deterministic quantities over the Boltzmann ensemble. This definition is general enough to be used in any biological context for which a ND GP map can be constructed. We use short RNA structures as a demonstration and validation of this approach.

### A. Neutral spaces and robustness

In a deterministic GP map the neutral set of a phenotype is the set of all genotypes that map to it. Its size as a fraction of genotype space is often referred to as the phenotypic frequency [13]. In the ND GP map we can define neutral sets in terms of the *average* frequency 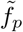 of a phenotype *p* over the Boltzmann probability distribution (as in refs [20, 21, 32]):

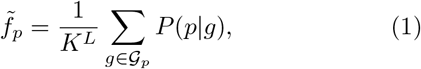

where 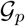 is the set of genotypes that contain the phenotype p in their Boltzmann ensemble and *P*(*p*|*g*) is the probability that genotype *g* gives phenotype *p*. *K^L^* is the total number of genotypes in genotype space, where *K* is the genotype alphabet size and *L* is the genotype sequence length. For RNA sequences of length *L*, the total number of genotypes in genotype space is 4^*L*^ since RNA has an alphabet of 4 nucleotides.

Robustness quantifies the insensitivity of a phenotype to genotypic mutations, and can be defined at the genotypic or phenotypic level [13]. In D GP maps, genotypic robustness *ρ_g_* corresponds to the fraction of genotypic point-mutation neighbours that leave the phenotype unchanged (also referred to as neutral neighbours) [13]:

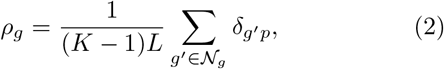

where 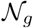 is the set of all point-mutation neighbours of *g, p* is the phenotype of *g* and *δ*_*g*′*p*_ equals 1 if the genotype *g*′ maps to phenotype *p*, and 0 otherwise. The normalisation (*K* – 1)*L* is the total number of point-mutation neighbours in genotype space. In RNA, the number of such neighbours for any genotype is 3*L*. If we want to generalise this definition to the non-deterministic case we need to consider the probability that *g* gives rise to phenotype *p* and the probability that genotypes in the mutational neighbourhoods of *g* also give rise to that phenotype. Thus, we define the non-deterministic genotypic robustness as:

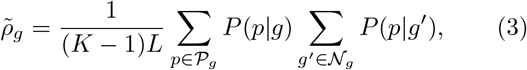

where 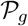 is the set of phenotypes *p* in the non-deterministic ensemble of genotype *g*, and 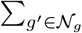 is the sum over all point-mutation neighbours *g*′ of *g*, which form the set 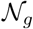. The maximum of 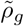 occurs when *g* only maps to one phenotype, and this phenotype is also the only phenotype for *g*′.

The deterministic phenotypic robustness *ρ_p_* is the genotypic robustness (Eq.2) averaged over all genotypes that have phenotype *p* [1, 13]:

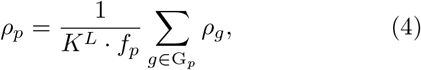

where G__p__ is the neutral set of *p* with size |*G_p_*| = *K^L^* · *f_p_*. For the non-deterministic phenotypic robustness we formulate an average of the non-deterministic robustness of a specific phenotype *p* over the non-deterministic neutral set size, resulting in the following definition:

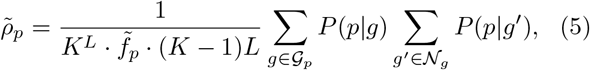

where the non-deterministic neutral set size is expressed using Eq.1, as 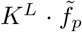. Equation 5 is analogous to the ND phenotype robustness definition recently proposed by Sappington and Mohanty [20].

### B. Evolvability

The capacity of a genotype or phenotype to produce phenotypic diversity is quantified by its ‘evolvability’ [13]. For the deterministic case, the genotypic evolvability of a given genotype *g* can be defined as the number of different structures found in its point-mutational neighbourhood [13]. The genotypic evolvability for the genotype *g* of phenotype *p* can therefore be written as:

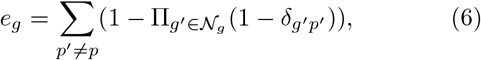

where as before 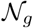 is the set of all point-mutation neighbours of *g*, and *δ*_*g*′*p*′_ equals 1 if the genotype *g*′ maps to the phenotype *p*′, and 0 otherwise. For the non-deterministic case the *δ*_*g*′*p*′_ is replaced by the probability *P*(*p*′|*g*′), resulting in an average of the number of different phenotypes in the neighbourhood of *g*. The average of each evolvability is weighted according to the probability that this phenotype is initially represented by *g*:

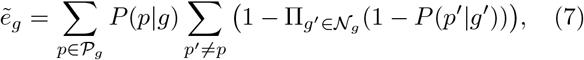

where the sum ∑_*p*′_≠_*p*_ is over all possible phenotypes *p*′ in the ND GP map except *p*.

The phenotypic evolvability of a phenotype *p* counts all different phenotypes in the mutational neighbourhood of *p* as a whole, or in other words, of all genotypes with phenotype *p* [13]. For the deterministic case, we can write this as:

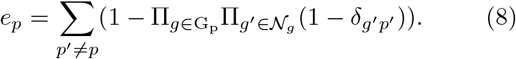

Note the additional product compared to Eq. 6 to make sure we are considering all *g*’s with *p*, represented by the neutral set G_*p*_. The non-deterministic phenotypic evolvability *ẽ_p_* represents the average number of accessible phenotypes from *p*, meaning the set of genotypes in 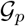. It can be formulated by replacing the *δ*_*g*′*p*′_ in Eq.8 with the product of probabilities *P*(*p*′|*g*′)*P*(*p|g*) and accounting for all genotypes in 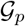:

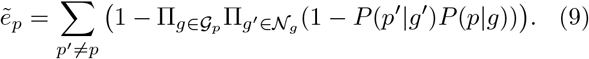

A simple GP map schematic such as in Fig.2 exemplifies the difference between the deterministic and non-deterministic case. The schematic shows the deterministic GP map as the map that assigns the most probable phenotype from the non-deterministic map for each genotype, which is analogous to how the MFE GP map of RNA relates to its ND GP map.

**FIG. 2.**
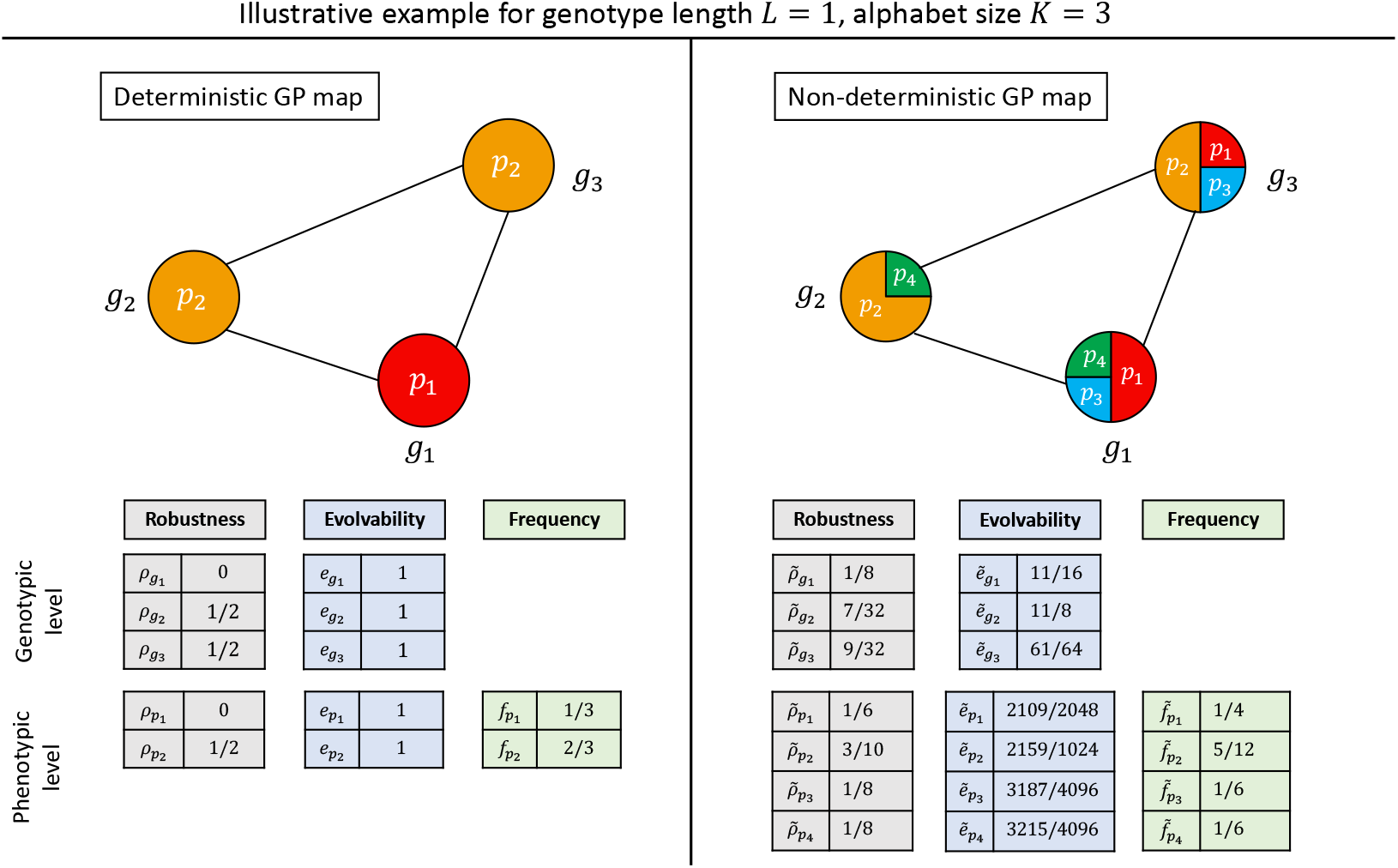
Schematic depiction of simple deterministic and non-deterministic GP maps, and associated properties. For an extremely simple genotype space of size 3 (genotype length *L* = 1 and alphabet size *K* = 3) we illustrate the difference between deterministic (D) and non-deterministic (ND) GP maps. The relative area that a phenotype occupies in a genotype node represents the probability that this phenotype arises from this genotype in the non-deterministic phenotype ensemble. Each node in the deterministic mapping is, like for the RNA MFE GP map, the most probable phenotype in the ND case. The robustness and evolvability of genotypes and phenotypes, as well as the phenotypic frequency are calculated for both the deterministic and non-deterministic case using Eq.1–9.

### C. Validation of non-deterministic quantities

We have defined the non-deterministic quantities in Eqs. 1, 3, 5, 7, and 9 to give us the averages of the deterministic quantities over a large number of realisations of the Boltzmann ensemble. To validate these non-deterministic definitions computationally we use short RNA, a biologically realistic computational GP map for molecular evolution. The ND GP maps for RNA of *L* = 12 and RNAshapes of *L* = 30 [31] are computationally constructed using the ViennaRNA package (see Methods). The validation consists of comparisons between the non-deterministic quantities for the ND GP map and the deterministic quantities averaged over a large sample (*N* = 500) of GP map realisations: each realisation is produced by drawing a sample from the Boltzmann distribution of each genotype, producing a single phenotype for each genotype in a given realisation. As can be seen in Fig.3A the non-deterministic definitions of robustness, evolvability and frequency match the averages of the deterministic quantities as expected, both at the genotypic and phenotypic levels. The phenotypic evolvability cannot be calculated with sampling methods as discussed in [33], therefore *e_p_* and *ẽ_p_* are not computed for RNAshapes30.

**FIG. 3.**
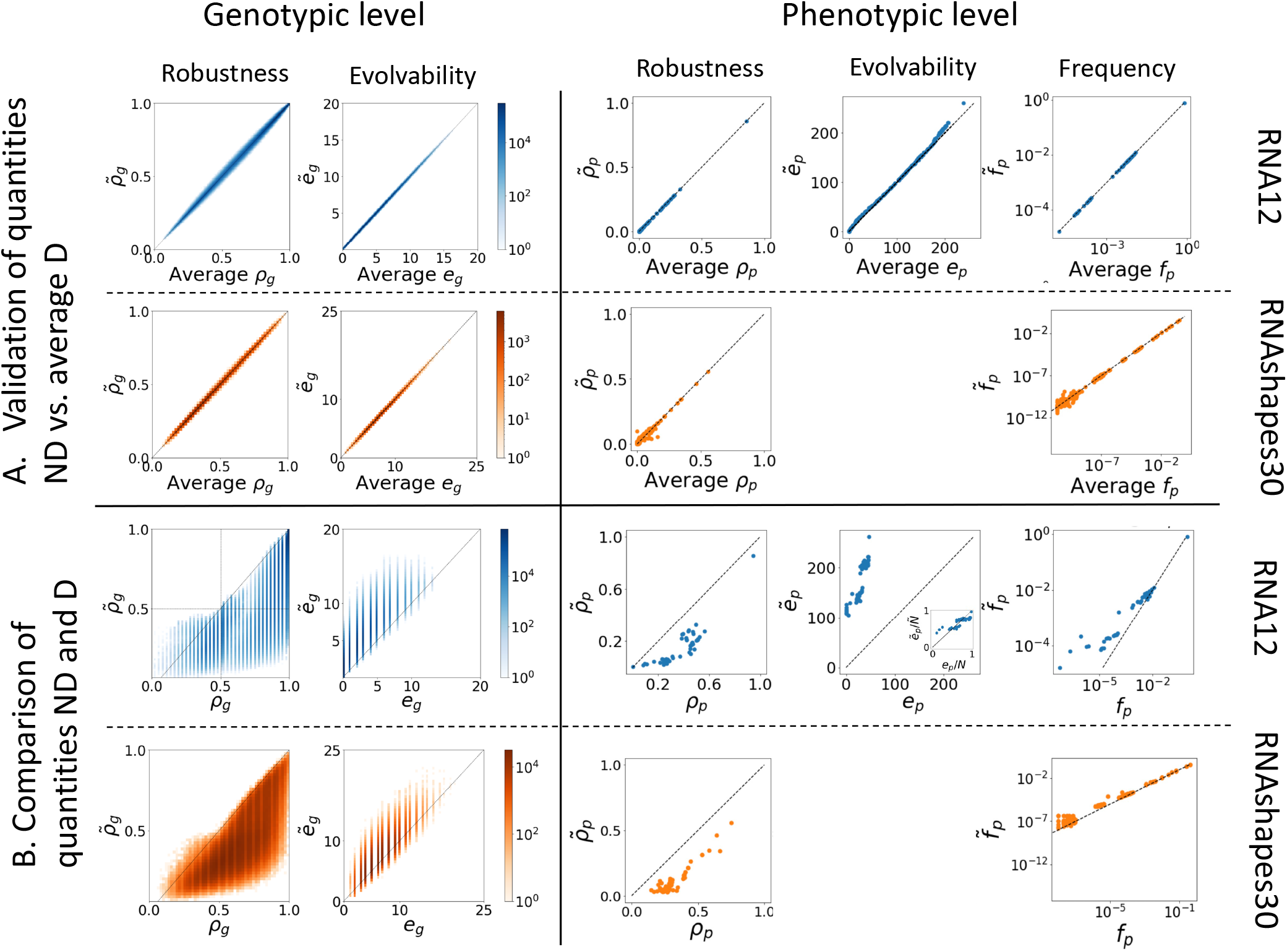
Non-deterministic framework validation and comparison. **A** The definitions of non-deterministic robustness 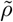, evolvability *ẽ*, and phenotypic frequency 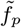 quantities are validated by comparing them to the averages of the equivalent deterministic quantities for RNA12 and RNAshapes30, across 500 samples of the ND GP map in both cases. **B**. Direct comparison between non-deterministic quantities on the ND GP map and the deterministic quantities on the MFE GP map. The phenotypic evolvability cannot be calculated through sampling methods [33], therefore we do not present evolvability plots for RNAShapes30. The phenotypic evolvability comparison for RNA12 includes an inset for normalised phenotypic evolvability *e_p_*/*N*, *ẽ_p_*/*Ñ*, which shows a very close correspondence of these two quantities (see full sized figure in Supplementary Information).

### D. Comparison of non-deterministic GP map properties to their MFE counterparts

We next compare the non-deterministic quantities in the ND GP map to their deterministic MFE counterparts. The latter are the GP map quantities that have been studied extensively in the RNA GP map literature (see Introduction). Fig.3B shows these comparisons.

At the genotypic level, Fig.3B shows similar behaviour for both RNA12 and RNAshapes30. We find that genotypes with high genotypic robustness in the MFE GP map tend to also have high robustness in the ND GP map. However, we find that in most cases the D genotypic robustness is higher than the ND genotypic robustness 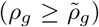, except for a low-density section of genotypes with 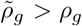 in the regime 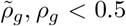. Similarly, genotypes with high genotypic evolvability in the MFE GP map tend to also have high evolvability in the ND GP map. However, genotypic evolvability is higher in the ND case (i.e. *ẽ_g_* ≥ *e_g_*) for the majority of genotypes. Therefore, we conclude that (most) genotypes in the ND GP maps of RNA secondary structure will on average be less robust and more evolvable than the same genotypes in the corresponding MFE GP map.

At the phenotypic level Fig.3B shows each corresponding quantity for the same phenotype. In the RNA12 ND GP map (energy range 15*k_B_T*, see Methods) there exist a total of *Ñ* = 271 phenotypes, while in the deterministic case there are only *N* = 48. Therefore, we can only compare the GP map quantities for the 48 phenotypes present in both maps. Given the result at the genotypic level, it is not surprising that phenotypic robustness is lower in the non-deterministic case than in the deterministic one. Phenotypic evolvability however shows a sharp increase, meaning that the same phenotypes are surrounded by many more phenotypes. However, this is a function of the increased total number of phenotypes found in the ND GP map, as shown by an inset plot comparing the *fractions* of all possible phenotypes that are discovered in the neighbourhood of a given phenotype *p*, which we will refer to as the normalised phenotypic evolvabilities *ẽ_p_/N* (for the ND GP map) and *e_p_/N* (for the D GP map). These normalised evolvabilities are similar in the MFE and ND GP maps (inset), but phenotypes at the low-evolvability end have slightly higher normalised evolvabilities in the ND case and thus a higher fraction of phenotypes is accessible from them. In all phenotypic plots for RNA 12 the outlier at the top right of the distribution represents the unfolded structure, which has a disproportionately large neutral space [14]. The phenotypic frequencies exhibit 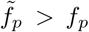 for the most part, particularly for low values of *f_p_* and 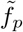. This means that low frequency phenotypes have larger neutral spaces in the ND GP map than in the MFE GP map, for both RNA12 and RNAshapes30.

### E. Structural properties of the ND GP map

Next we determine whether the structural properties that have been observed across several GP maps, including the RNA MFE GP map (see Introduction) also hold in the ND GP map. First, the so-called ‘bias’ observed in many GP maps describes the highly skewed distribution of phenotypic frequencies [5]. In most GP maps the vast majority of genotypes maps to a small number of phenotypes, and the remainder maps to many different phenotypes. This property is also found in the non-deterministic RNA GP maps because the phenotypic frequency values, 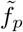, differ by orders of magnitude, as shown in Fig.3 (as found in refs [20, 21, 32]). Second, many GP maps, including the RNA MFE GP map, exhibit a negative correlation of genotypic evolvability and robustness and a positive correlation of phenotypic evolvability and robustness [13]. The negative correlation in the genotypic case is expected as the neighbourhood of an individual genotype cannot contain many instances of the same phenotype and also instances of many other phenotypes. A trade-off is therefore inevitable. The positive correlation at the phenotypic level however is a consequence of the shape and size of neutral networks, as well as the organisation of information into biological sequences [13, 34, 35]. These relationships also hold in the case of the ND GP map, as can be seen in Fig.4 for RNA12. At the genotypic level, the expected MFE GP map negative correlation is present (Pearson correlation coefficient of −0.73) and a similar negative correlation can be observed in the ND GP map (Pearson correlation coefficient of −0.85). At the phenotypic level the expected positive correlation is observed for the MFE GP map (Pearson correlation coefficient of 0.81). The ND GP map also shows a positive correlation (Pearson correlation coefficient of 0.61), but the points are somewhat differently distributed. If we distinguish MFE phenotypes (highlighted in red, and also plotted separately in the inset) among all phenotypes in the ND GP map, we recover a distribution that mirrors the MFE case more closely. The phenotypes that only appear in the ND GP map form a long tail with very low robustnesses but a wide range of evolvability values.

**FIG. 4.**
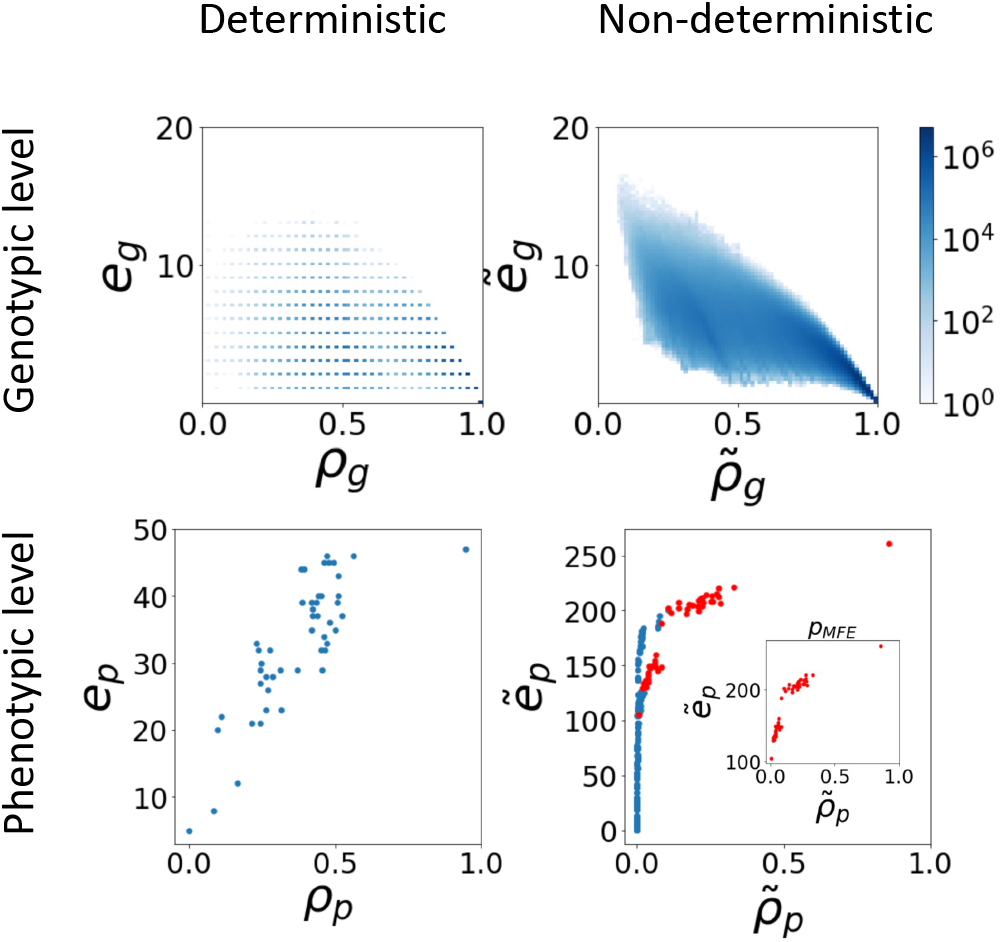
RNA12 robustness versus evolvability. At the genotypic level evolvability and robustness exhibit negative correlations, with a Pearson correlation coefficient of −0.73 in the MFE GP map, and −0.85 in the ND GP map. At the phenotypic level the correlation between evolvability and robustness is positive with Pearson correlation coefficients 0.81 (MFE GP map) and 0.61 (ND GP map). The structures that appear in both the MFE and ND GP map are highlighted in red in the ND phenotypic evolvability/robustness plot.

In Fig.5 the RNAshapes30 display a similar relationship between evolvability and robustness at the genotypic level, in the form of negative correlations for both the MFE and ND GP maps (Pearson coefficient of −0.56 for the MFE GP map and −0.64 for the ND GP map). Again, because the phenotypic evolvability cannot be calculated with sampling methods [33], we do not study the correlations at the phenotypic level for RNAshapes30. These results indicate that the commonly observed correlations between evolvability and robustness are also present in the ND GP map.

**FIG. 5.**
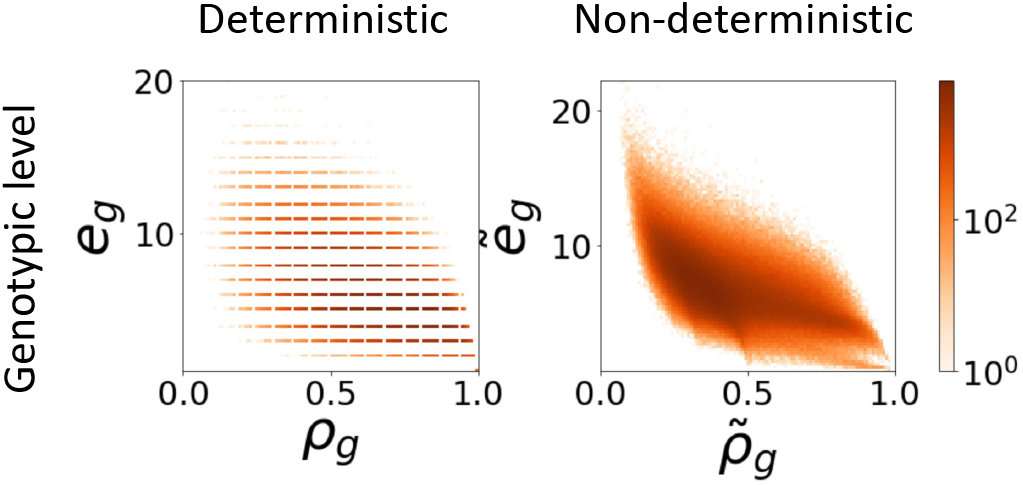
RNAshapes30 robustness versus evolvability. The MFE GP map shows a negative correlation between the genotypic robustness *ρ_g_* and genotypic evolvability *e_g_*, with a Pearson coefficient of −0.56. A negative correlation is also observed in the ND GP map with Pearson coefficient of −0.64.

Many GP maps, including the RNA MFE GP map, display neutral correlations [14]. One might expect these to also be found in the ND GP map, due to plastogenetic congruence. This concept, introduced by Ancel and Fontana [15] relates mutational neighbourhood of a structure to its Boltzmann distribution (see Supplementary Information). To verify that the ND GP map also contains neutral correlations we verify whether 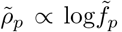 holds, as has been found for D GP maps [4, 10, 14]. Fig. 6 shows the relationship between these two quantities, and verifies the presence of correlations as for most phenotypes the robustness 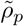 is significantly larger than the null model representing a random distribution of pheno-types with 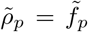, as previously found for the D and recently also the ND GP maps [14, 20]. We also observe, in both the RNA12 and RNAshapes30 ND GP maps, the presence of the recently observed biphasic robustness scaling [20]: For higher-frequency phenotypes, neutral correlations are present 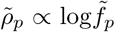 but suppressed compared to the MFE GP map, which follows from the lower phenotypic robustness found in Fig. 3; in lower-frequency phenotypes, the correlations disappear and the relationship becomes analog to the null model 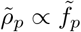 [20].

**FIG. 6.**
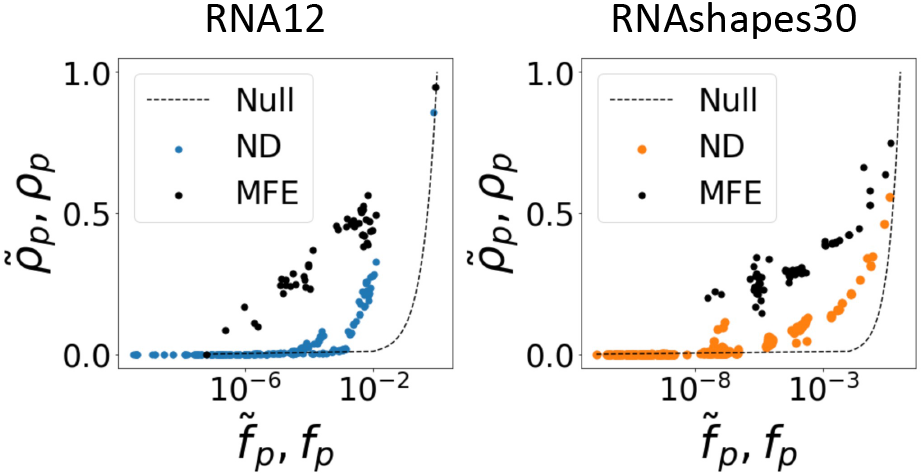
Presence of neutral correlations: phenotypic robustness versus phenotypic frequency for RNA12 and RNAshapes30. Neutral correlations are still present in the ND GP maps for high phenotype frequencies, 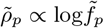. The robustnesses of low-frequency phenotypes equal those of the null model, 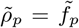. In the MFE GP maps the neutral correlations are present for all frequencies, 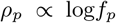, as previously observed [14].

## III. CONCLUSION

The study of GP maps and their structural properties provides an important perspective on evolutionary processes that complements the widely-studied concept of the fitness landscape [28, 36]. For example, a determinant feature that has driven the evolution of natural functional RNAs is the accessibility of the phenotypes in the GP map which depends on the neutral networks and their distribution [5]. This suggests that the GP map structure and its implications for variation can be just as important as selection pressures [5], as new phenotypes need to be accessible in order to be selected. Robustness, evolvability and neutral spaces have become definitions in the general framework to quantify the relationship between the genotype and phenotype in GP maps, and have been key to the exploration of the quasi-universal structural properties seen in GP maps in different biological contexts [1, 37]. Our contribution is a generalisation of this framework to non-deterministic GP maps, which represent a form of phenotypic plasticity [28]. The study of plasticity, its effects on evolution, and the evolution of plasticity itself has been a field of interest in evolutionary biology since the late nineteenth century [22, 24, 38]. Plasticity can arise as a result of interactions with the environment or through a variety of physical, chemical and biological mechanisms[39]. In molecular evolution, plasticity plays a fundamental role since molecules are exposed to thermal fluctuations and produce plastic phenotypes in the form of a statistical distribution of folded structures. The study of plastic or non-deterministic GP maps therefore provides a more realistic picture of variation and its impact on molecular evolution.The results discussed in this paper provide a foundation for further research on non-deterministic GP maps and future work should apply the quantities defined here to other examples of ND GP maps, for example proteins or GRNs. A particularly interesting avenue to explore would be how these non-deterministic quantities are applied to evolutionary dynamics.

## IV. METHODS

### A. RNA12

The ND GP map is constructed by computing the Boltzmann ensemble of suboptimal structures and their probabilities for each RNA sequence of length *L*. For the case of RNA *L* = 12 the folds are calculated for every sequence through ViennaRNA with all parameters set to default values (e.g. the temperature *T* = 37°C), using the ViennaRNA suboptimal function [7] and the Boltzmann probabilities of these are obtained using the partition function. The energy range for the suboptimals is 15*k_B_T* as in [40] to be consistent with the RNAshapes data. The final ND GP map is constructed by mapping each genotype sequence to its ensemble of structures in the energy range (including unfolded structure), as well as their respective normalized probabilities.

### B. RNAshapes for length *L* = 30

The RNAshapes calculations rely on the implementation in ref [40], which uses the ViennaRNA package (version 2.4.14) [7]. This is faster than the original RNAshapes program for short sequences and thus allows us to work with larger samples of sequences and structures. We use the same parameters as in ref [40] (shape level 2, temperature *T* = 37°C, energy range 15*k_B_T*, sequence length *L* = 30, no isolated base pairs). Unlike ref [40], however, we compute the deterministic map by taking the most frequent structure in the Boltzmann ensemble as the deterministic folded structure, without additional requirements on its relative frequency.

Since there are 4^30^ ≈ 10^18^ genotypes of sequence length L = 30, we have to rely on sampling: we simply work with 10^6^ randomly generated sequences (10^5^ for the average case, which is computationally more expensive due to the necessary repetitions). For the plots showing genotype robustness and evolvability, we simply plot one value for each sequence in the sample - thus, not all possible genotypes are in the sample, but we obtain an overview of common genotypic robustness and evolvability values for a large random sample of genotypes. For the phenotypic robustness estimates, we simply approximate each sum over all sequences by a sum over our sequence sample. This approach was tested for RNA12, where we have exact data as a reference (shown in the Supplementary Information, section S1.1). To investigate whether the chosen sequence sample is large enough, we analysed how robust our results are to sub-sampling (shown in the Supplementary Information, section S1.2). Phenotypic evolvabilities are not estimated since they cannot be inferred reliably from samples with current techniques [33]. Phenotypic frequencies were also estimated from a random sequence sample. However, phenotypic frequency calculations are faster than robustness calculations because they do not require us to examine the mutational neighbours of each sequence, and so we used a larger sample of 10^8^ sequences. Note that our reliance on sampling methods means that quantities for low-frequency phenotypes, which do not appear in the sample, cannot be estimated.

## ACKNOWLEDGMENTS

PGG was funded by a “La Caixa” Foundation postgraduate scholarship and NSM acknowledges funding from the Issachar Fund.

## Supplementary Information

### S1 Validation of sampling methods for the phenotype robustness calculations for the *L* = 30 RNAshapes map

#### S1.1 Test case: sampling for the RNA12 map

**Figure S1:**
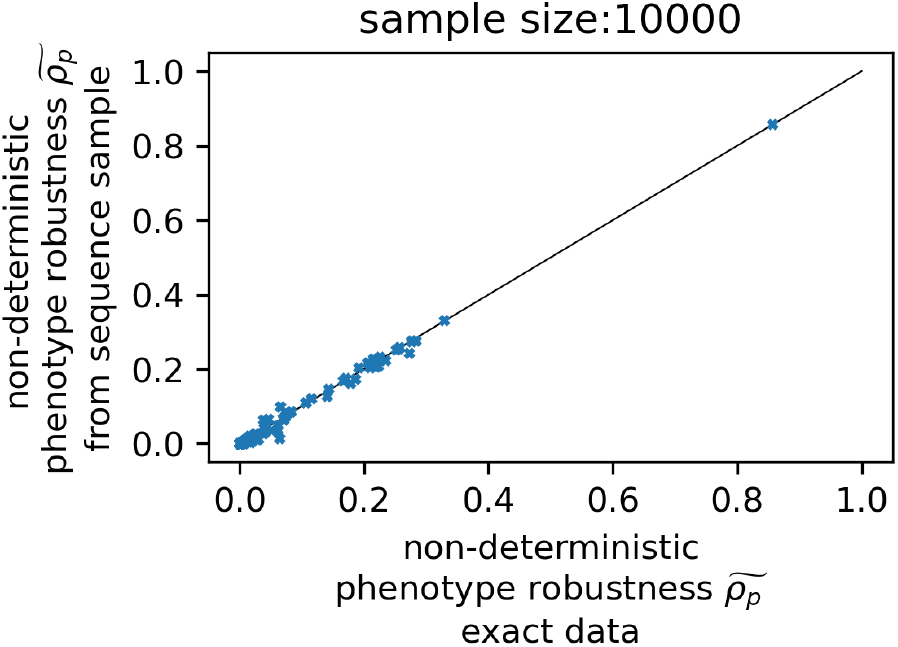
Non-deterministic phenotype robustness estimates for the RNA12 map: *the non-deterministic phenotype robustness estimate for each structure is computed from the full sum over all genotypes (x-axis) and estimated based on a partial sum over* 10^4^ *randomly selected genotypes (y-axis). We find a good agreement, even though the fraction of all genotypes that are sampled* (10^4^/4^12^ ≈6 × 10^-4^) *is much smaller than several phenotypic frequencies*.

For the RNA12 map, we can sum over all genotypes exhaustively and thus compute the non-deterministic phenotype robustness exactly and without sampling. Thus, we use this map as a test case, to check if the non-deterministic phenotype robustness can be inferred accurately from a sample. This is shown in Fig S1: here the sample-based estimates are plotted against the exact values of the non-deterministic phenotype robustness for all structures of length *L* = 12. We find that there is excellent agreement between the two methods, even though the sample size is small: only 10^4^ sequences are used in the sample. This means that the sample size is smaller than the phenotypic frequencies of several structures and thus some structures are unlikely to appear as MFE structures in the sample, but only as suboptimal structures. Despite this, the non-deterministic phenotype robustness can be estimated from the sample with reasonable accuracy, indicating that a random sequence sample is an appropriate choice of sample for the phenotypic frequency.

Thus, a random sequence sample does not introduce systematic errors in the non-deterministic phenotype robustness estimates for RNA12, and we assume that the same applies for longer sequence lengths and the RNAshapes ensembles. The same also holds for the deterministic map (see the supplementary material of ref [1] for a discussion).

#### S1.2 RNAshapes data: sensitivity to sample size

**Figure S2:**
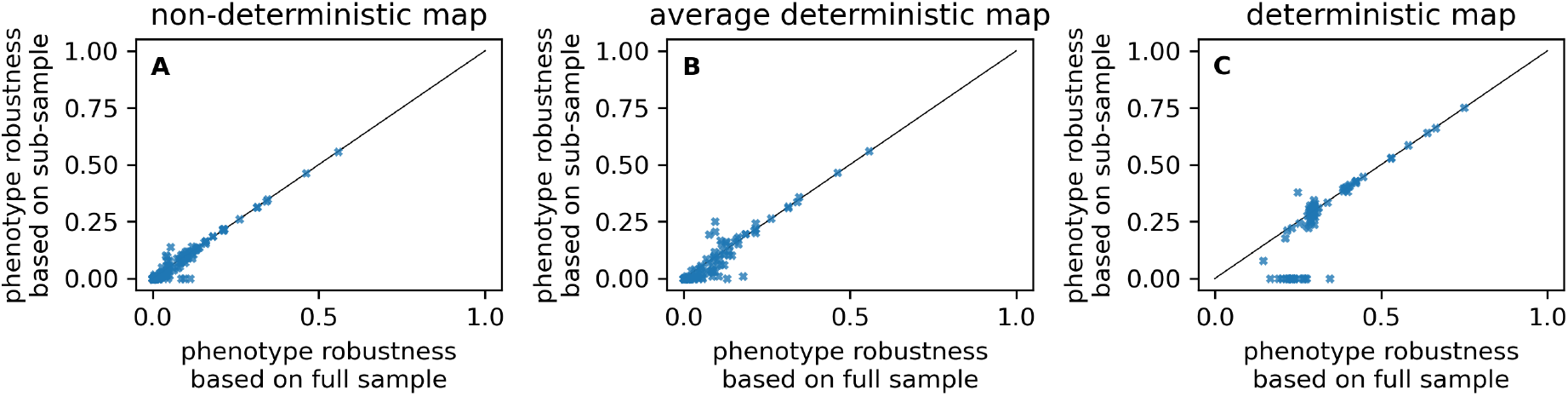
Effect of sub-sampling on the phenotype robustness values for the *L* = 30 RNAshapes data: *in order to test, if our sample size is sufficiently large, we also compute phenotypic robustness values for a sub-sample consisting of* 10% *of the sequences in the full sample and plot these values against the values derived from the full sample. This analysis is shown for A) the non-deterministic GP map, B) the average deterministic map, C) the deterministic map*.

Here, we test if the sample size in our RNAshapes30 calculations is sufficiently large. To test this, we repeat our sample-based phenotype robustness calculations for a subsample consisting of 10% of the sequences in the full sample (Fig S2). While sampling errors are clearly present in our analysis, especially for phenotypes with low robustness in the deterministic map, there is a high correlation between the values calculated from the full sample and those calculated from a sub-sample. This indicates that sampling errors are present, but are small compared to the true differences in phenotype robustness.

### S2 Normalised evolvability

The evolvability quantifies the accessibility of phenotypes at the genotypic or phenotypic level [2]. The quantity can be normalised by the total amount of different phenotypes in the full GP map. This normalised evolvability gives a proportion of accessible phenotypes, which becomes useful if trying to compare phenotypic evolvability between two different GP maps. In Fig S3 the normalised phenotypic evolvability for the RNA12 MFE GP map and ND GP map is compared, showing that out of the phenotypes that have a low proportion of accessible phenotypes in their neutral space, these proportions become larger in the non-deterministic case compared to the deterministic one. The unfolded structure corresponds to the highest evolvability value and remains of similar value in both maps.

**Figure S3:**
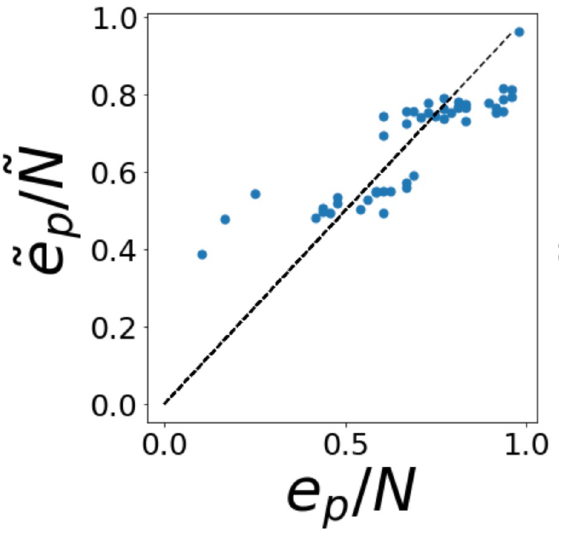
Normalised evolvability comparison between the MFE D GP map and ND-GP map for RNA12: *The phenotypic evolvability is normalised by the total number of different phenotypes in the GP map, for the case of RNA12 MFE D GP map this number N is 48 and for the ND GP map Ñ is 271.*

### S3 Plastogenetic congruence

The influential paper by Ancel and Fontana [3] explored the structural properties of the plastic RNA GP map. They observed a correlation between an RNA genotype’s MFE structure and the suboptimal structures of its 1-mutation neighbours, which they called plastogenetic congruence [3]. In that time, GP maps were not being systematically analysed in their complete sequence-structure mapping because of computational limitations, and so the concept was previously tested based on samples [4]. Now, this is possible and plastogenetic congruence can be checked for in the full RNA ND GP map. In this paper we have used the full sequence-structure RNA12 ND-GP map for energy range 15*k_B_T*. To observe plastogenetic congruence we expect a positive correlation between the probability of the MFE structure *P*(*p_mfe_*|*g*) vs. the genotypic robustness of that genotype *ρ_g_* in the MFE GP map, as shown in Fig.S4. The plot has a positive correlation with Pearson *r* = 0.64. We conclude that our RNA12 ND GP map has the expected structural feature of plastogenetic congruence.

**Figure S4:**
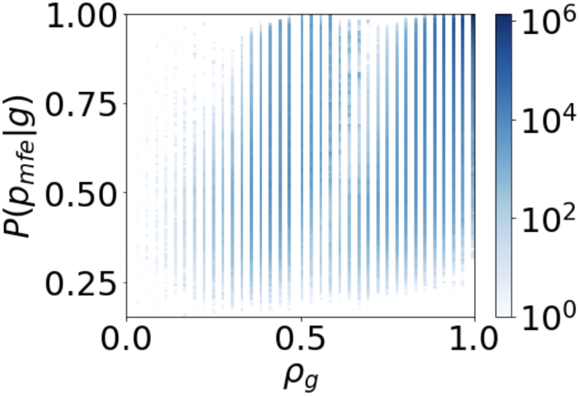
Plastogenetic congruence in the computationally constructed RNA12 ND GP map: *There is a positive correlations (Pearson r = 0.64) between the probability of the MFE structure in the Boltzmann ensemble, and the genotype’s robustness [3].*

